# Modeling relaxation experiments with a mechanistic model of gene expression

**DOI:** 10.1101/2024.04.04.588028

**Authors:** Maxime Estavoyer, Marion Dufeu, Grégoire Ranson, Sylvain Lefort, Thibault Voeltzel, Véronique Maguer-Satta, Olivier Gandrillon, Thomas Lepoutre

## Abstract

**Background:** In the present work, we aimed at modeling a relaxation experiment which consists in selecting a subfraction of a cell population and observing the speed at which the entire initial distribution for a given marker is reconstituted.

**Methods:** For this we first proposed a modification of a previously published mechanistic two-state model of gene expression to which we added a state-dependent proliferation term. This results in a system of two partial differential equations. Under the assumption of a linear proliferation rate, we could derive the asymptotic profile of the solutions of this model.

**Results:** In order to confront our model with experimental data, we generated a relaxation experiment of the CD34 antigen on the surface of TF1-BA cells, starting either from the highest or the lowest CD34 expression levels. We observed in both cases that after approximately 25 days the distribution of CD34 returns to its initial stationary state. Numerical simulations, based on parameter values estimated from the dataset, have shown that the model solutions closely align with the experimental data from the relaxation experiments.

**Conclusion:** Altogether our results strongly support the notion that cells should be seen and modeled as probabilistic dynamical systems.

## Background

Cells are neither machines [1] nor simple information processing devices [2, 3]. Their specific complexity sometimes led to the idea that they should be treated differently that classical physico-chemical systems [4]. Nevertheless like all living systems cells are rooted within a physico-chemical reality which they can not escape. We therefore argue that cells should be seen and modelled as probabilistic dynamical systems.

One obvious sign that cells should indeed be seen as such lie in the possibility to perform so-called “relaxation” experiments. This consists in selecting a subfraction of a cell population (potentially down to one cell) and observing the speed at which the entire initial distribution for a given marker is reconstituted. Such relaxation experiments have already been published and analyzed on various cells and antigens. Arguably the very first report to do so analyzed the distribution of the Sca1 antigen (Stem Cell Antigen 1) at the surface of EML cells, a multipotent mouse haematopoietic cell line. It was shown that it took more than 9 days before the three fractions (most Sca-1 negative, most Sca-1 positive and a central fraction) regenerated Sca-1 histograms similar to that of the parental (unsorted) population [5]. The authors proposed a phenomenological model which point toward discrete transitions in a dynamical system exhibiting multistability to quantitatively predict the relaxation dynamics of the sorted subpopulations [5]. For this they assumed the existence of two stable states, one of low and one of high Sca1 expression. Proliferation was assumed to be equal in both states.

Other studies have adopted a somewhat different approach with the knock-in of fluorescent reporter genes under the control of endogenous promoters [6, 7]. The first targeted promoter was that of Nanog in murine embryonic stem cells [6], an other gene classically considered as a stemness marker. Similarly to [5], the authors demonstrated that, although being in a Nanog low of in a Nanog high state is not biologically equivalent in term of fate, the transition between these two states can be adequately modelled using a fully probabilistic model, simulated using a Stochastic Simulation Algorithm [8].

The second targeted promoter was that of Tenascin-C in NIH 3T3 mouse fibroblasts [7]. In that case, the authors first proposed a phenomenological 2-states model, which proved to not correctly capture their data. They then turned toward a Langevin type stochastic differential equation to model the relaxation process. This led to an accurate prediction of the rates at which different phenotypes will arise from an isolated subpopulation of cells [7]. In contrast with [5], the authors assumed that each state had its own proliferation rate.

In the present work, we aimed at using a previously published mechanistic model of gene expression [9] to which we will add a stemness-dependant proliferation term, to fit relaxation data obtained by examining CD34 expression at the surface of TF1-BA cells.

## Methods

### Mathematical model

#### Case without proliferation

Throughout this work, we will use the classical two-state model (Figure 1; see [9] and references therein), a refinement of the pioneer model introduced by [10].

**Fig. 1.**
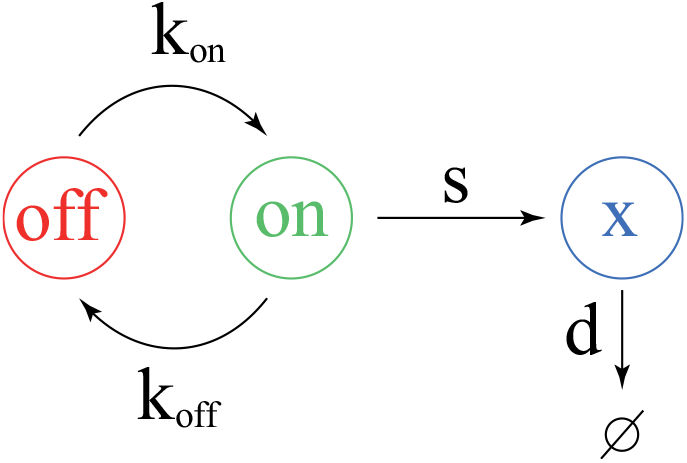
The 2-state model of gene expression. The gene opens with a *k*_on_ rate and closes with a *k*_off_ rate. Similarly to [11] we only consider protein (*x*) production (with an *s* rate) and degradation (with a *d* rate)

This is the simplest model that accounts well for the specific nature of single-cell omics data (non-poisonnian [12], well fitted by Gamma distributions [13] and displaying a high proportion of zero counts [14]). More refined models with any number of possible gene configuration have been described [15] but their mathematical complexity makes them cumbersome to use for our purpose.

It is important to stress here that this is a mechanistic model, that differs from the phenomenological 2-states model described upper. Such models only considered a low and a high *χ* state, without describing the protein production process. Importantly here stochasticity is described at the core of the modelling and does not need to be introduced as a additional term in the model. We recently proposed a piecewise deterministic Markov process (PDMP) version of that model which rigorously approximates the original molecular model [9]. Furthermore, a moment-based method has been proposed for estimating parameter values from a given experimental distribution assumed to arise from the functioning of a 2-states model [11].

The master equation of the process in the absence of proliferation reads

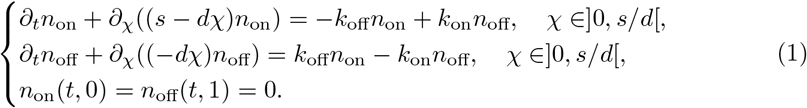

We define *X*_max_ as the maximum value for the quantity of CD34 in a cell. Scaling the space by *X*_max_ allows us to consider the following system

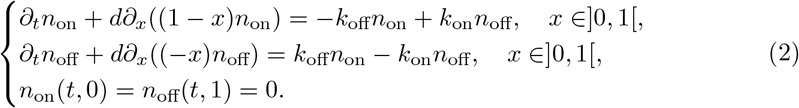

with *n*_on_(*t, x*) being the number of cells with a promoter in the on state at time t, with a (scaled) CD34 level *x* and *n*_off_(*t, x*) being the number of cells with a promoter in the off state. The total number of cells, denoted as *n*(*t, x*), is given by *n*_on_(*t, x*) + *n*_off_(*t, x*). This is the quantity we considered to be measured.

##### Steady state of the model

The system is mass preserving and it converges to a steady state *N*_on,off_ which is characterized by

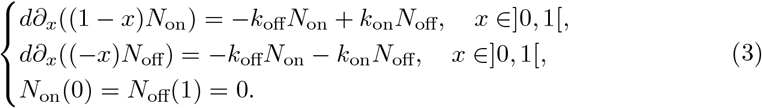

And the solution if nonnegative. An interesting feature of this system is the fact that we have an explicit solution. Indeed, summing up the equations, we get

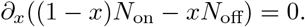

Therefore, this quantity is constant on]0, 1[. Using the boundary condition, we can see that 0 is the only admissible constant. Therefore, in this precise case, we have necessarily

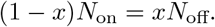

Injecting in the equation we get

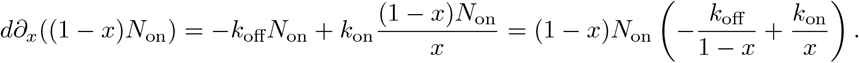

From this, we get

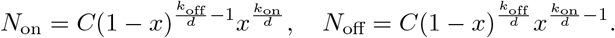

and quite remarkably, we have

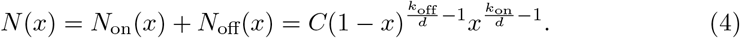

If we choose 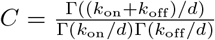 we normalize this to 1 and get a *β* law *B*(*k*_on_, *k*_off_), so that we end up with

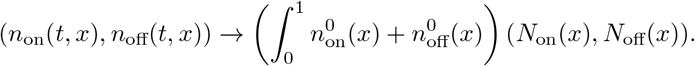

#### Case with proliferation

HSCs mostly reside in a quiescent state, although they can occasionally divide during homeostasis [16]. We therefore will consider CD34^+^ TF1-BA cells as immature slowly proliferating cells and CD34^−^ TF1-BA cells as mature highly proliferating cells. Therefore the proliferation rate will depend on the *x* variable representing the CD34 content but not on the on*/*off status.

Moreover, we consider that cells keep their on*/*off status during a division. This is in line with the demonstration of a memory of transcriptional activity in mammalian cells [17, 18].

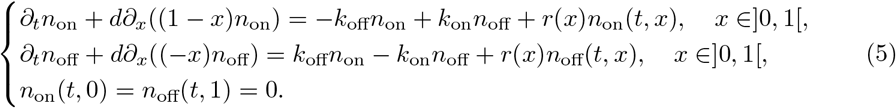

Since the system is structurally non-conservative, it makes no sense to look for steady state here. However, one can investigate for stable exponential profile, that is to look for positive solutions with shape

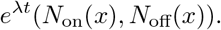

Such solution satisfy the system

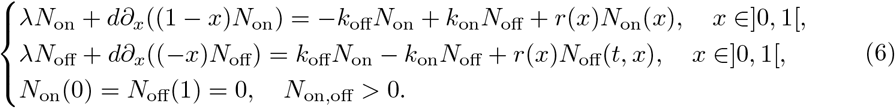

In the sequel, we will focus on the normalized representant so that we will assume

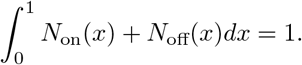

We also introduce the adjoint eigenprofile

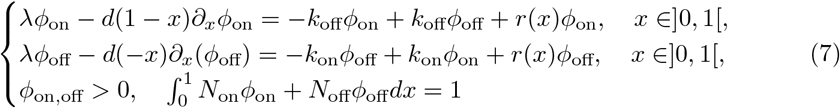

We emphasize in particular the following property, for any initial condition of the system, one has

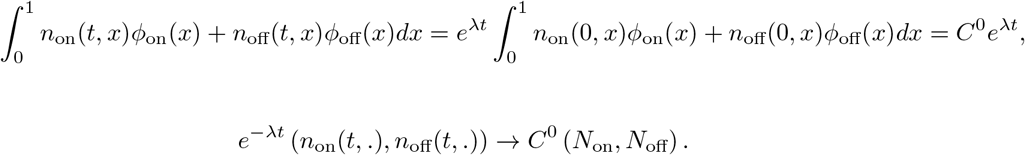

For more details on this, we refer to [19] for an introduction to eigenvectors in this context. In particular, thanks to our normalization, the triplet (*λ, N, ϕ*) is uniquely defined. Note that in the conservative case (*r* = 0), *λ* = 0, *N* is given by the renormalized steady state and the adjoint eigenvector is simply the constant vector (1, 1). Note also that this guarantees that for any initial data, we have

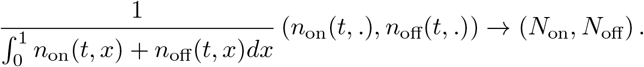

And regarding the observations of the steady profile in figure 2, our normalized asymptotic profile should be *N* : *x* ↦ *N*_on_(*x*) + *N*_off_(*x*).

**Fig. 2.**
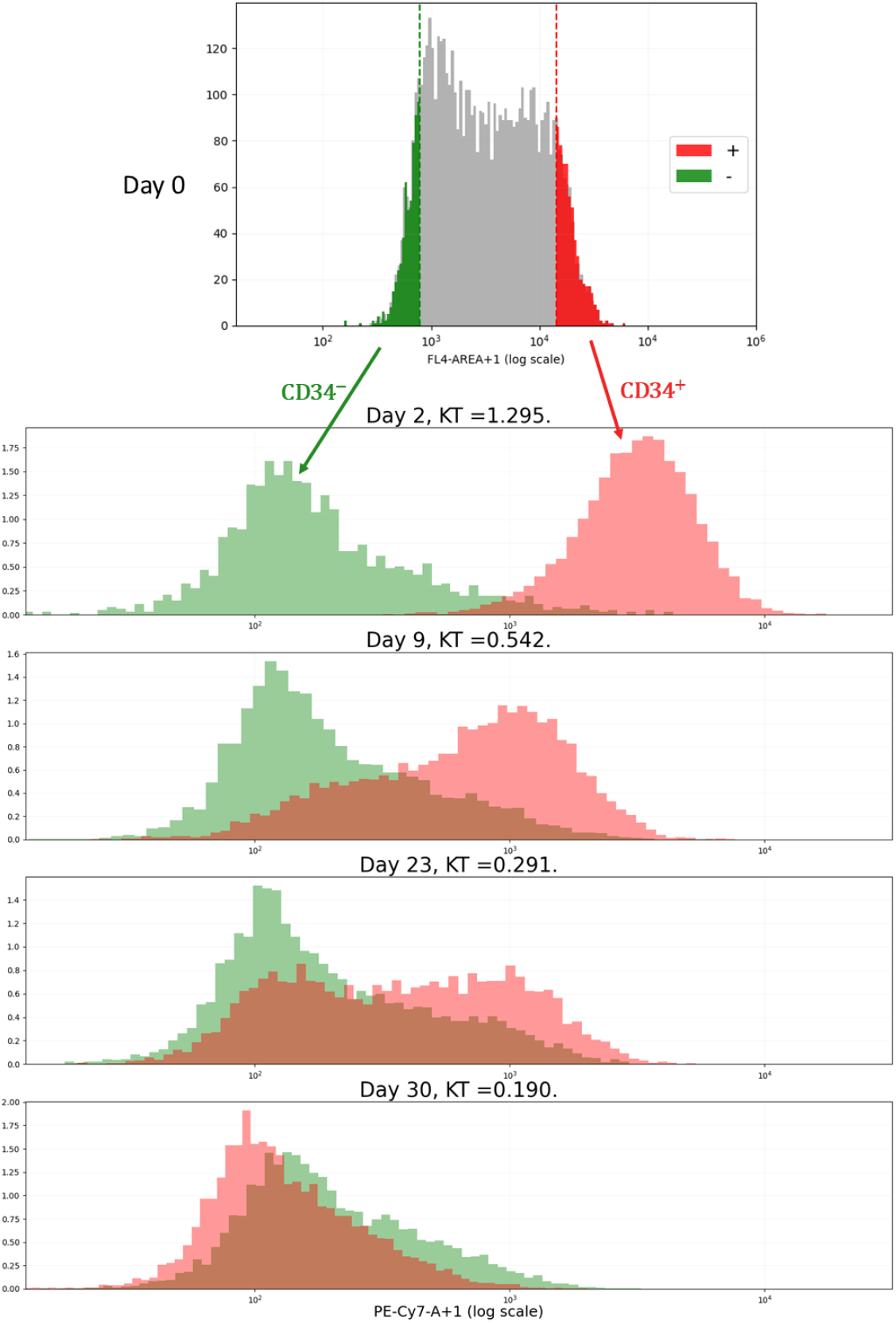
The relaxation experiment. TF1-BA cells well labelled with an anti-CD34 antibody and FACS-sorted. The 10 percent most CD34 positive and the 10 percent most CD34 negative cells were sorted, grown in culture for the indicated period of time, where the distribution of cell-surface CD34 expression was assessed. KT: the modified Kantorovich-Rubinstein distance, defined by the equation (14), between the two distributions [23].

We assume that, for the initial dimensional system, the proliferation rate is linear, *i*.*e*. 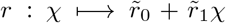. Scaling again the space by *X*_max_, the proliferation term becomes, *r* : *x → r*_0_ +*r*_1_*x* with 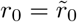 and 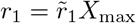. We assume that the constant proliferation rate is positive, *r*_0_ *>* 0. Conversely, to model the fact that CD34^−^ cells divide more frequently than CD34^+^ cells, we assume that the linear proliferation term, *r*_1_, is negative. However, to preserve the positivity of the proliferation rate, the constant *r*_1_ must satisfy the following condition, *r*_1_ ∈ [− *r*_0_, 0].

We show in the Results section that, under this proliferation assumption, it is theoretically possible to derive the normalized asymptotic profiles (*N*_on_, *N*_off_).

### The biological setting

#### Relaxation experiments

Chronic Myeloid Leukemia (CML) is a myeloproliferative disorder arising at the hematopoietic stem cell (HSC) level. It is associated with the recurrent chromosomal (Philadelphia) translocation t(9;22)(q34;q11) which leads to the oncogenic chimeric gene that fuses Bcr and Abl genes and results in the expression of a constitutively active unique tyrosine kinase named BCR-ABL [20].

Véronique Maguer-Satta’s group has developed the TF1-BA cell line, a unique model of immature human hematopoietic cells (TF1) transformed with BCR-ABL by lentiviral infection. This model was shown to recapitulate early alterations following the BCR-ABL oncogene appearance as identified using primary samples of CML patients at diagnosis and in chronic phase [21].

We decided to analyze the relaxation dynamics for the CD34 antigen at the surface of those TF1-BA cells (Figure 2). CD34 is a transmembrane phosphoglycoprotein which is predominantly regarded as a marker of Haematopoietic Stem Cells (HSCs) [22]. We reasoned that CD34 surface expression could therefore be seen as a proxy for stemness of our TF1-BA cells. Interestingly, one observes a relaxation in both directions: CD34^−^ cells are regenerated from CD34^+^ cells, as biologists would expect, but one also see that CD34^+^ cells are regenerated from CD34^−^ cells, establishing that stemness is not a fixed quality but the result of an underlying dynamical system as previously shown in other cellular systems ([5, 6]; see upper)

#### Data processing

Two types of data were collected on days 2, 5, 9, 13, 19, 23, 26 and 30 : cell counts and fluorescence distribution. The cell counts allow us to quantify proliferation whereas the fluorescence measure the distribution of CD34 expression.

##### Gating

As usual for cytometric data, we initiate the analysis with a gating step. We use SSC-H and FSC-H data to distinguish viable and debris cells. FSC (Forward Scatter) data are generally assimilated to the size of the cells analyzed, and correspond to the light scattered along the laser path. SSC (Side Scattered) data, on the other hand, are usually linked to the granularity and correspond to the light scattered at a 90-degree angle. The “H” stands for Height and is one component of this type of data. Cell debris are characterized by relatively low size and high granularity relative to their size.

To select viable cells, we plot the values of SSC-H along those of FSC-H. An example of such a graph is provided in Figure 3 using data from day 2 of the CD34^+^ cell experiment. Visually, viable cells can be identified as the cluster of points with high FSC-H and SSC-H values. Using the “FlowCal” python package [24], we draw an ellipse as a filter to select only these viable cells. At the bottom of Figure 3, we represent fluorescence data with and without the gating phase (*Ungated* and *Gated*, respectively). Note that fluorescence distribution is only slightly affected by the removal of debris cells.

**Fig. 3.**
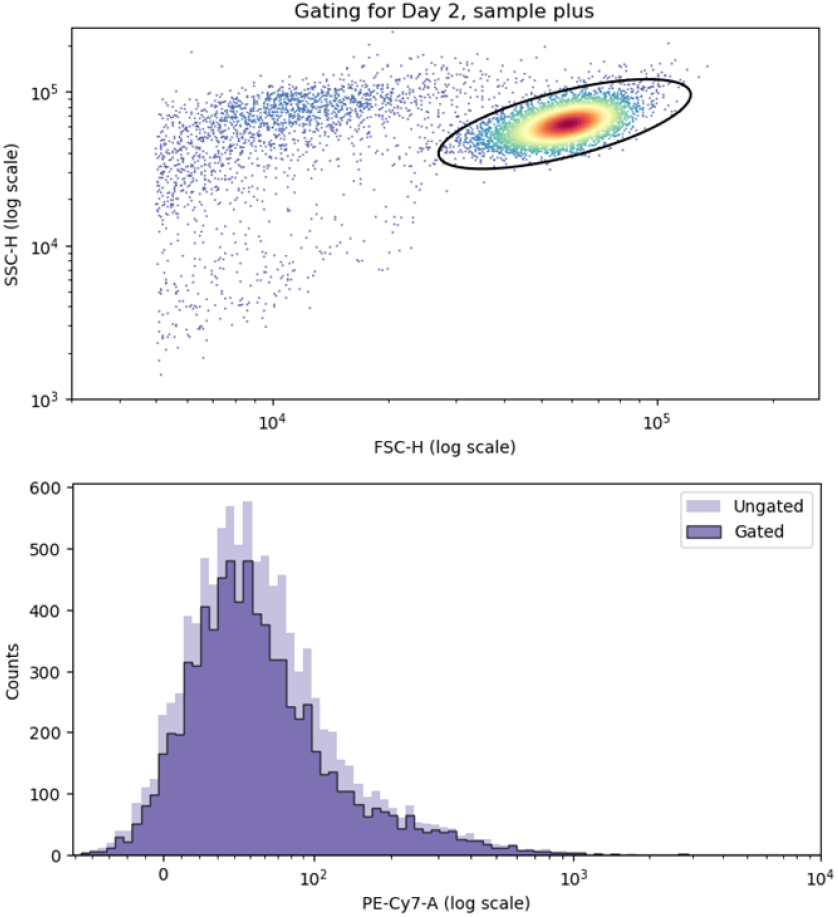
Example of flow cytometry gating. *Top*. Example of SSC-H along FSC-H plot for raw data from the plus subpopulation on day 2. As the data contain a high proportion of debris cells, we select only those viable cells lying within the black ellipse. *Bottom*. Fluorescence data before gating (*Ungated*) and after gating (*Gated*). For the figures and the ellipse, we used the python package “FlowCal” [24].

##### Shifting

Even after gating, some cells exhibit a negative fluorescence level, which is inconsistent as these values are intended to represent the amount of proteins in each cell. To avoid this problem, we added a shifting step. This step occurs immediately after the gating process and consists in subtracting the minimum value of each distribution (for each sub-population and for each day) from all the values, bringing the minimum to 0. This transformation, once again, does not distort the fluorescence distribution.

### Numerical simulations

#### Linking data to mathematical model

The cell counts are interpreted as snapshots of the total population 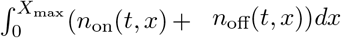. The fluorescence distribution is considered as a sample from the distribution 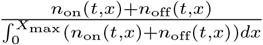. As we have no information on the repartition on*/*off for the initial data, we apply the following rule : for *t*_0_, starting of our simulation (DAY 2), we choose the repartition to be the same as in the steady distribution *N*. More precisely, we fix the proportion with the equation

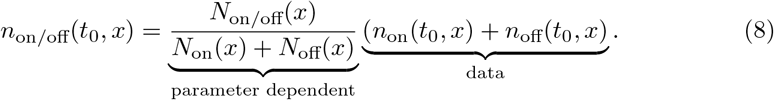

#### Numerical scheme

For the equations (5), we use an explicit upwind scheme. Setting *a*_on_ : *x* ↦ *d*(1 − *x*) and *a*_off_ : *x* ↦ −*dx*, the scheme is given by

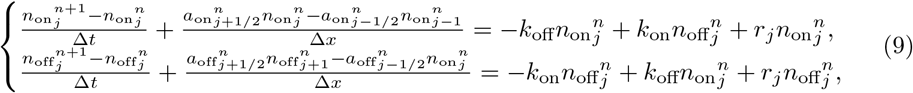

with 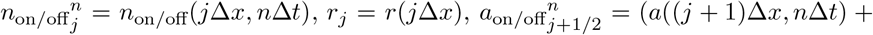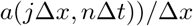 and 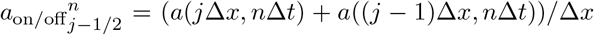. In the Results section, figure A1 illustrates a comparison between the result of the numerical scheme and the theoretical asymptotic profile of equations (5).

#### Estimation of the distance to the data

To calibrate the parameter values of our system, we use our experimental data. Initially, in order to estimate the exponential growth rate of cells, we perform a linear regression analysis on the temporal evolution data of the cell count. To determine the values of other system parameters, we seek values that make our numerical results as close as possible to the experimental data. To characterize this notion of closeness between our numerical results and the data, we introduce the Kantorovich-Rubinstein distance. Given two probability distribution *p*_1_, *p*_2_ on ℝ _+_, we define their cumulative distribution function *P*_i_ (*x*) = *Pr*(*X > x*, under distribution 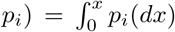. Using these functions we can define the Kantorovich Rubinstein (also known as Wasserstein) distance by

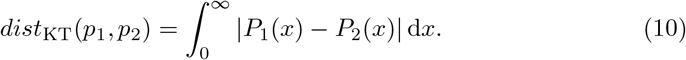

In our specific case, we want to compare at each step the (normalized) distribution generated by the model at time *t*_*i*_ (hereby denoted as *model*(*t*_*i*_, *dx*) with cumulative distribution *M* (*t*_*i*_,.)) and the corresponding distribution of the data at time *t*_*i*_ denoted as *data*(*t*_*i*_, *dx*)with cumulative distribution *D*(*t*_*i*_,.). We would therefore compute

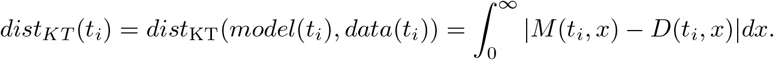

Note that in our case the integral is in fact taken on the finite interval [0, 1] for scaled variables.

Considering the distribution profile of the data, we prefer to study this distance on a logarithmic scale. We therefore make the following change of variables *y* = log(*xX*_max_+ 1), and we define the modified Kantorovich-Rubinstein distance as follows

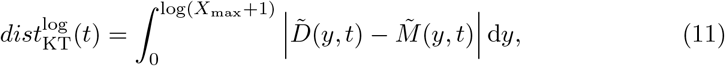

with 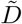 and 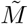 the two cumulative distribution functions, defined below, in the new logarithmic scale. Note that, following this change of variable, this “distance” can be greater than 1.

We are looking for a function 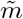 that satisfies the following relation, for all *b* ∈ [0, log(1 + *X*_*max*_)]

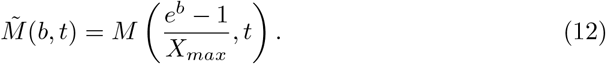

In particular, the link between the corresponding densities is immediately given by

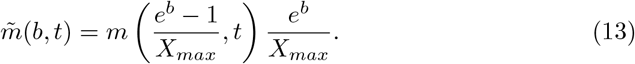

The space [0, 1] is discretized uniformly with *J* + 1 points, and this sequence is denoted (*x*_*j*_)_*j*_.

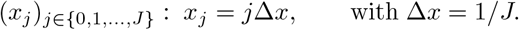

We also define the sequences (*y*_*j*_)_*j*∈*{*0,1,…,*J}*_ and (*l*_*j*_)_*j*∈*{*0,1,…,*J*−1*}*_ as follows

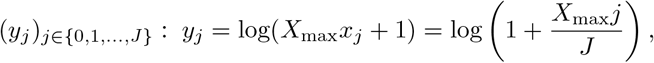

and

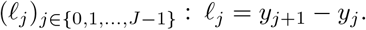

Consequently, the estimator of the cumulative distribution function *M* is given by

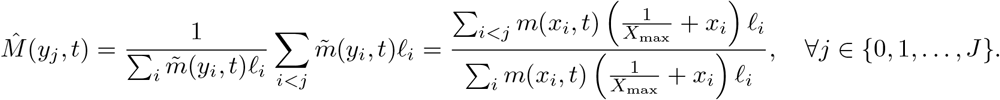

where we have used (13) to estimate 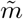. We need to renormalize to ensure we are comparing probability distributions.

The estimator of the cumulative distribution function *D* is

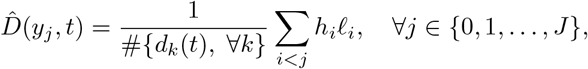

with *h*_*j*_(*t*) = #{*d*_*i*_ : log(*d*_*i*_(*t*) + 1) ∈ [*y*_*j*−1_, *y*_*j*_]}, where the operator # corresponds to the cardinal of a set and the data *d*_*i*_ correspond to the fluorescence data obtained after data processing. These data, *d*_*i*_, are real numbers between 0 and *X*_max_. The term # *d*_*k*_(*t*), ∀*k*} corresponds to the number of cells present in the data on day *t* after the gating operation.

Therefore, the distance between the experimental data and the mathematical model is as follows

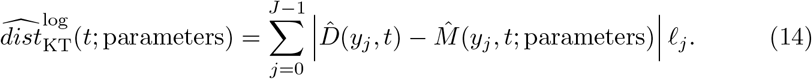

To calibrate the parameters of our model, we will minimize the sum of the modified Kantorovich-Rubinstein distance for the different days at our disposal and for the two experiments. The distance associated with CD34^+^ data is denoted 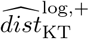, while that associated with CD34^−^ data is denoted 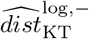. We also introduce the distance, denoted 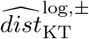, corresponding to the sum of these two distances. Mathematically, the optimization problem is given by the following formula

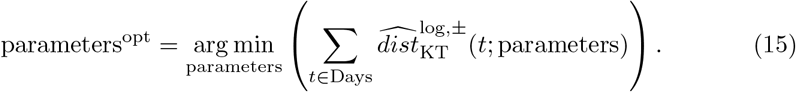

The results of this optimization work are detailed in the Results section.

## Results

### Mathematical Analysis – derivation of explicit normalized asymptotic profile (*N*_on_, *N*_off_)

Under the assumption of a linear proliferation rate *r*(*x*) = *r*_0_ + *r*_1_*x*, we obtain the following result

#### Theorem 1.

*Assume r*(*x*) = *r*_0_ + *r*_1_*x. Define the matrix M by*

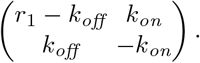

*Denote s*(*M*) *the largest eigenvalue of M, then the left and right eigenvector*

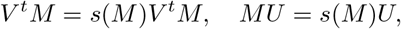

*can be chosen positive. Moreover, the solution to* (6) *is given by*

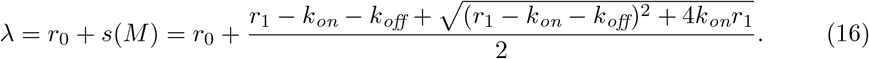

*And*

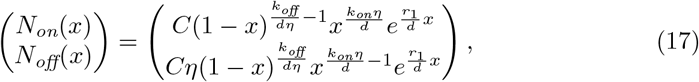

*with C an arbitrary positive constant and η given by η* = 1 + *s*(*M*)*/k*_*on*_.

*In particular, we have*

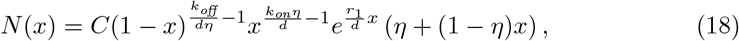

*Proof*. We notice that the existence and uniqueness (up to a multiplicative factor) of the triplet *s*(*M*), *V, U* is a straightforward consequence of the classical Perron Frobenius theorem which applies here because the off-diagonal entries of M are positive [25]. We introduce the ration 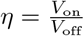 and notice that by construction, we have

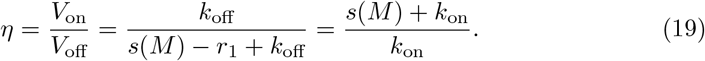

Then, consider the system satisfied by 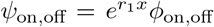, where *ϕ*_*on/off*_ is the solution of (7). We get

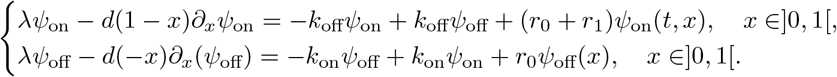

This system can be summarized as

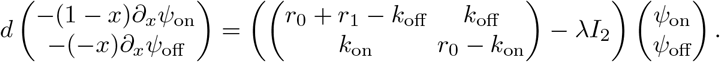

We recognize the matrix *r*_0_ + *M* ^*t*^. Therefore, we have a solution independent on *x Ψ*_on,off_ = *V*_on,off_ and *λ* = *r*_0_ + *s*(*M*) and 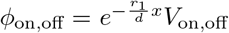. Similarly, if we denote 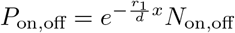, we obtain

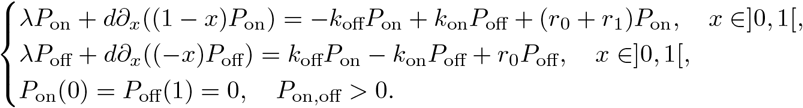

This can be condensed into

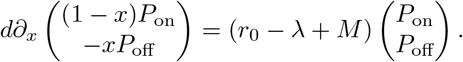

We can now proceed as for the conservative case and notice that since *V* ^*t*^(*r*_0_ − *λ* − *M*) = 0 multiplying the equation by *V*.

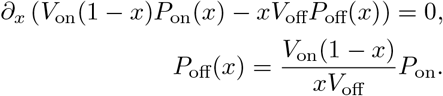

After the appropriate substitution, we obtain

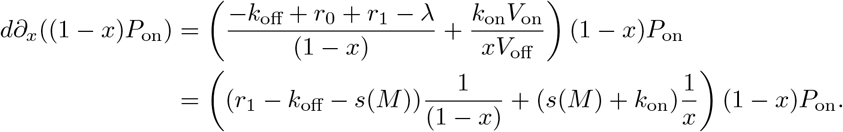

This leads to, for a suitable renormalization constant *C >* 0,

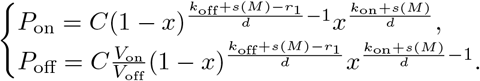

We introduce then the notation 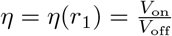 and go back the *N* variables to write

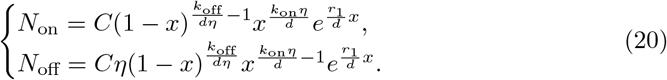

In particular, we have

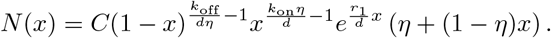

Going back to the definition of *η* we notice

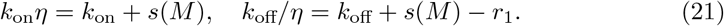

□

### Simulation Analysis

#### Estimation of exponential growth rate

Using experimental data on cell numbers for different days, we estimate the exponential growth rate *λ*. For both relaxation experiments, we perform a linear regression of the natural logarithm of the number of cells. In the case of CD34^+^ cell relaxation, the linear regression line is given by the slope *λ*^+^ ≈ 0.418, and for CD34^−^ cells, the slope is *λ*^−^ ≈ 0.422. We estimate the parameter *λ* by the average of these two slopes, *λ* ≈ 0.42. Figure 4 shows that the estimate of the exponential growth rate is in good agreement with the experimental data.

**Fig. 4.**
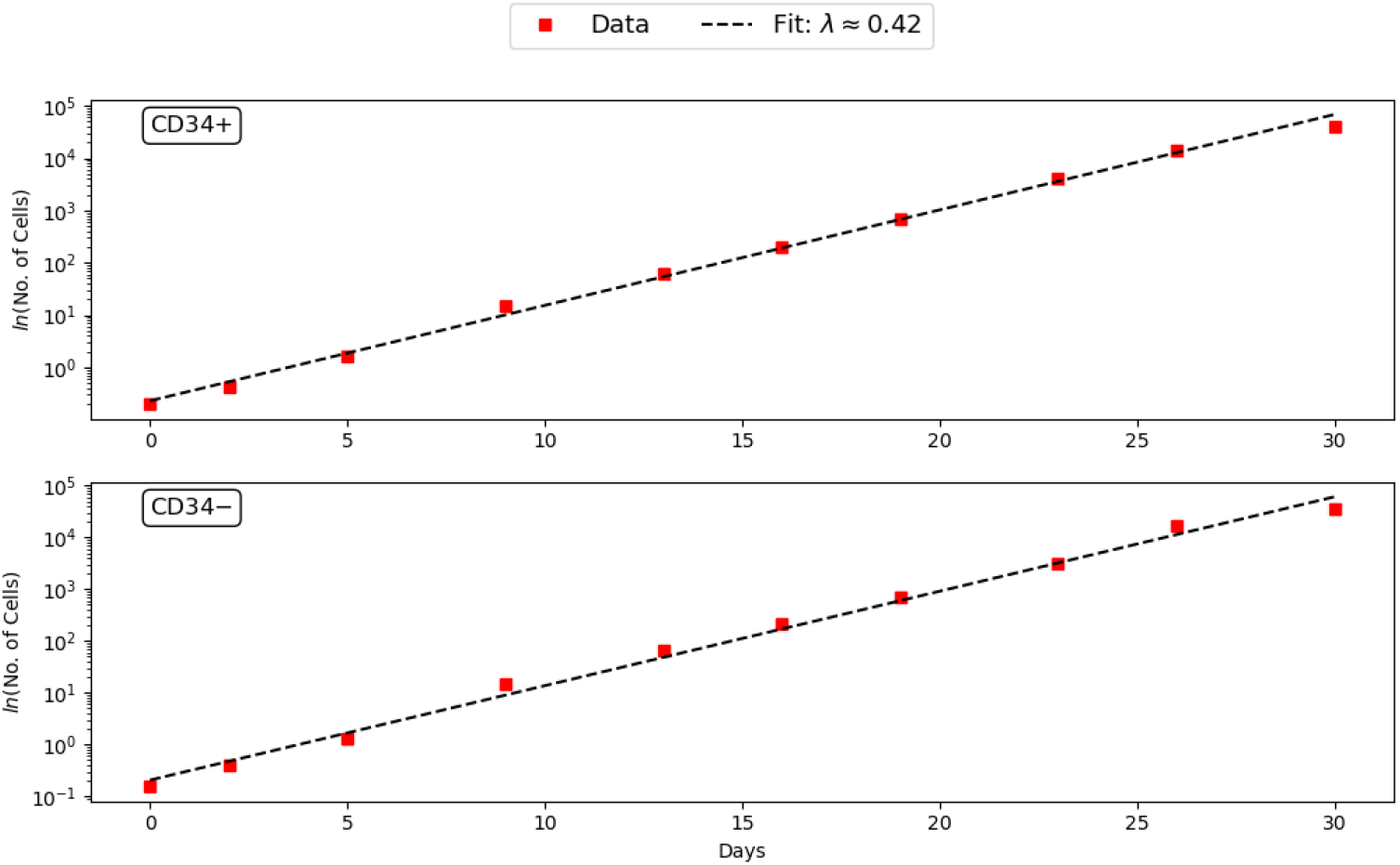
Estimation of the exponential growth rate *λ*. The red squares correspond to the number of cells (log scale) at different times for the relaxation experiments: *Top*. CD34^+^, *Bottom*. CD34^−^. The dotted line in black illustrates the optimal fit of the experimental data. The average of the slopes of the linear regressions minimizing the two experiments is given by the slope *λ* ≈ 0.42.

#### Calibration of parameters

A study of the maximum for each day and each experiment of the “PE-Cy7-A” fluorescence data reveals an *X*_max_ close to 20,000. To reduce the numerical complexity of the 5-parameter optimization (*r*_0_, *r*_1_, *k*_on_, *k*_off_, *d*), we will use the estimate of *λ* to reduce our optimization problem to just 4 parameters. Indeed, using the theoretical relationship (16) and the previous estimate of *λ*, we can define the parameter *r*_0_ as a function of the other model parameters,

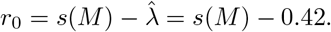

For the estimation of the other parameters *r*_1_, *k*_on_, *k*_off_ and *d*, we use the modified Kantorovich-Rubinstein distance minimization strategy, presented in the Method section and given by the following formula

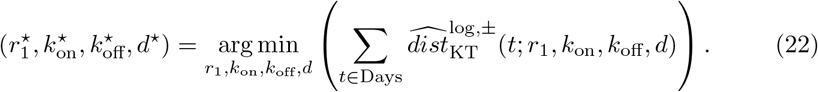

To determine numerically this minimum, we calculate the modified Kantorovich-Rubinstein distance for the two relaxation experiments, for each point on a large grid of parameter values. We then adjust our grid to obtain the location of the minimum of the sum of two distances. This method gives us the following four parameter values 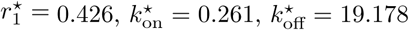 and *d*^⋆^ = 0.21. Following this optimization strategy, the optimal choice for the function *r* is to choose *r*_1_ such that *r*(*x*) = *r*_0_ × (1 − *x*). Nevertheless, we will see in the next section that the choice of *r*_1_ is not decisive for a good fit between the model result and the experimental data.

Finally, using the relation, *s* = *dX*_max_, we can calculate the value of the synthesis rate, *s* = 4214. All parameter value estimates are given in table 1.

**Table 1.** Estimated parameter values.

| Parameter | Value | Units | Description | Estimation Method |
| --- | --- | --- | --- | --- |
| $X_{\max}$ | $2 \times 10^4$ | proteins | Maximum value for the quantity of CD34 in a cell | Data-driven selection |
| $r_0$ | 0.426 | $\text{h}^{-1}$ | Constant proliferation rate | Estimating the proliferation rate $\lambda$ and using the relation (16) |
| $r_1$ | -0.426 | $\text{h}^{-1}$ | Linear proliferation rate | KT distance minimization (22) |
| $k_{\text{on}}$ | 0.261 | $\text{h}^{-1}$ | Rate at which the gene/promoter is turned "on" | KT distance minimization (22) |
| $k_{\text{off}}$ | 19.178 | $\text{h}^{-1}$ | Rate at which the gene/promoter is turned "off" | KT distance minimization (22) |
| $d$ | 0.21 | $\text{h}^{-1}$ | Degradation rate | KT distance minimization (22) |
| $s$ | 4214 | proteins.$\text{h}^{-1}$ | Synthesis rate of proteins when gene is on | Relation: $s = dX_{\max}$ |

#### Profile likelihood

To investigate the robustness of our parameter estimates and the significance of each parameter in minimizing the modified Kantorovich-Rubinstein distance, we employ an approach analogous to the *profile likelihood* concept [26, 27] in the context of our optimization problem.

First, we examine the influence of the parameter *d*. Let *d* be fixed, we calculate, in the same way, the triplet of parameters (*r*_1_, *k*_on_, *k*_off_) that minimizes the modified Kantorovich-Rubinstein distance under the fixed *d* constraint. These optimal parameters are, therefore, functions dependent on *d*, denoted as 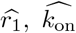, and 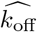.

Mathematically, they are defined by the following relation

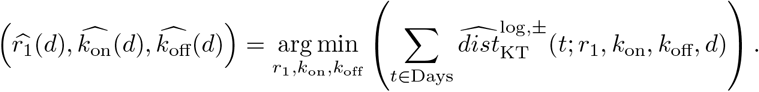

Once these functions have been calculated, we can determine the modified Kantorovich-Rubinstein distance associated with them, denoted by *S*_*d*_ and defined by the following equality,

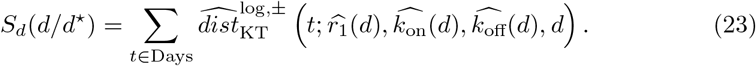

For the argument of the function, we choose *d/d*^⋆^, to study the distance associated with the relative variation of the optimal parameter. By definition of the function *S*_*d*_, it reaches its minimum at *d* = 1, corresponding to *d* = *d*^⋆^. Similarly, we can define the functions 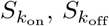, and 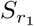.

In figure 5.A, we plotted the 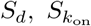 and 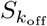 functions. The impact of the relative variation of the two transition rates around the optimal value, on the modified Kantorovich-Rubinstein distance is quite similar. For the degradation rate, *d*, we note that a fine estimate of this is crucial to obtain good accuracy between the data and the mathematical model.

**Fig. 5.**
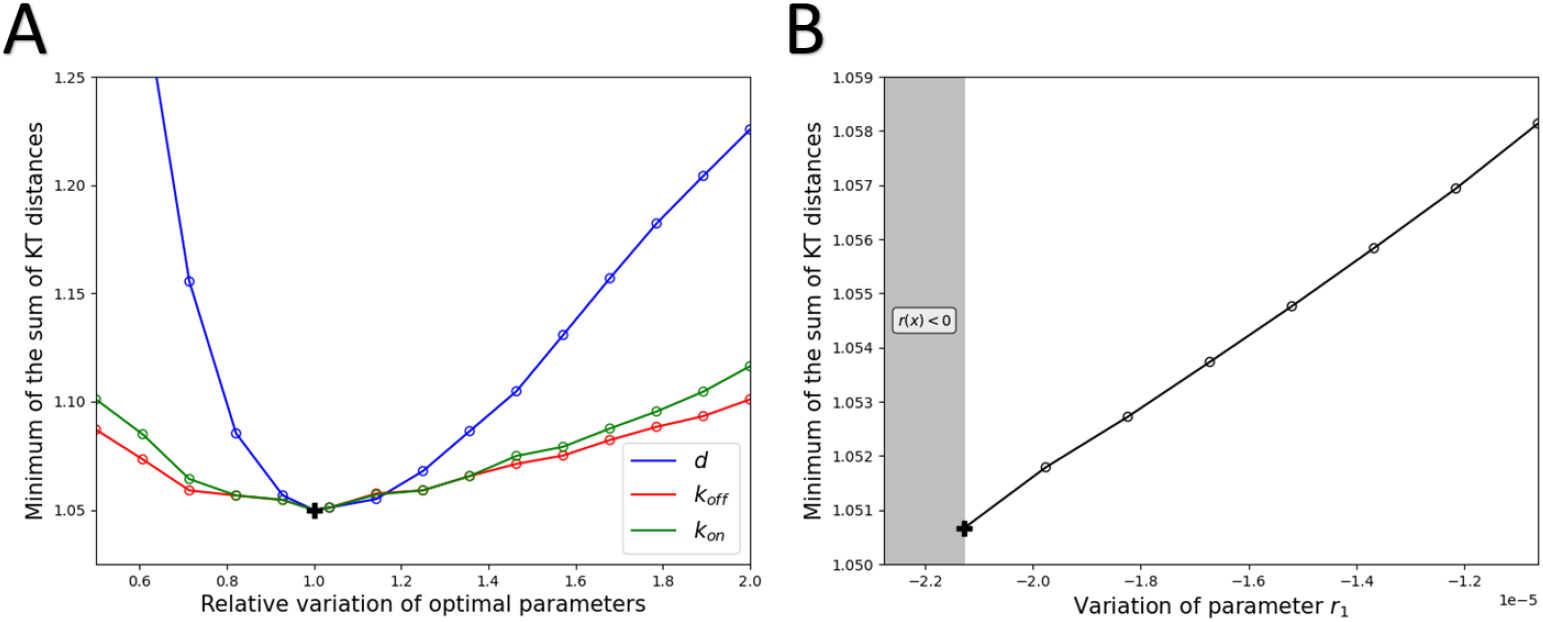
Likelihood profiles for for *k*_on_, *k*_off_ and *d* in A and for *r*_1_ in B. A. The blue curve represents the function *S*_*d*_, the red curve 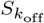 and the green curve 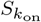. The function *S*_*d*_ is introduced into equation (23). B. The grey area corresponds to the range of parameter values for *r*_1_ such that the function *r* is non-positive. Compared with the other parameters, variation in the *r*_1_ parameter has a small impact on the minimum Kantorovich-Rubinstein distance.

Conversely, the parameter *r*_1_ has a minor impact on the minimum of the modified Kantorovich-Rubinstein distance. Specifically, when *r*_1_ deviates from its optimal value, new optimal parameter values emerge, resulting in distances very close to the optimal distance. This result is illustrated in Figure 5.B.

#### Comparison between model and experimental data

In figure 6 we compared data from relaxation experiments with the results of our model for the parameter values presented in table 1.

**Fig. 6.**
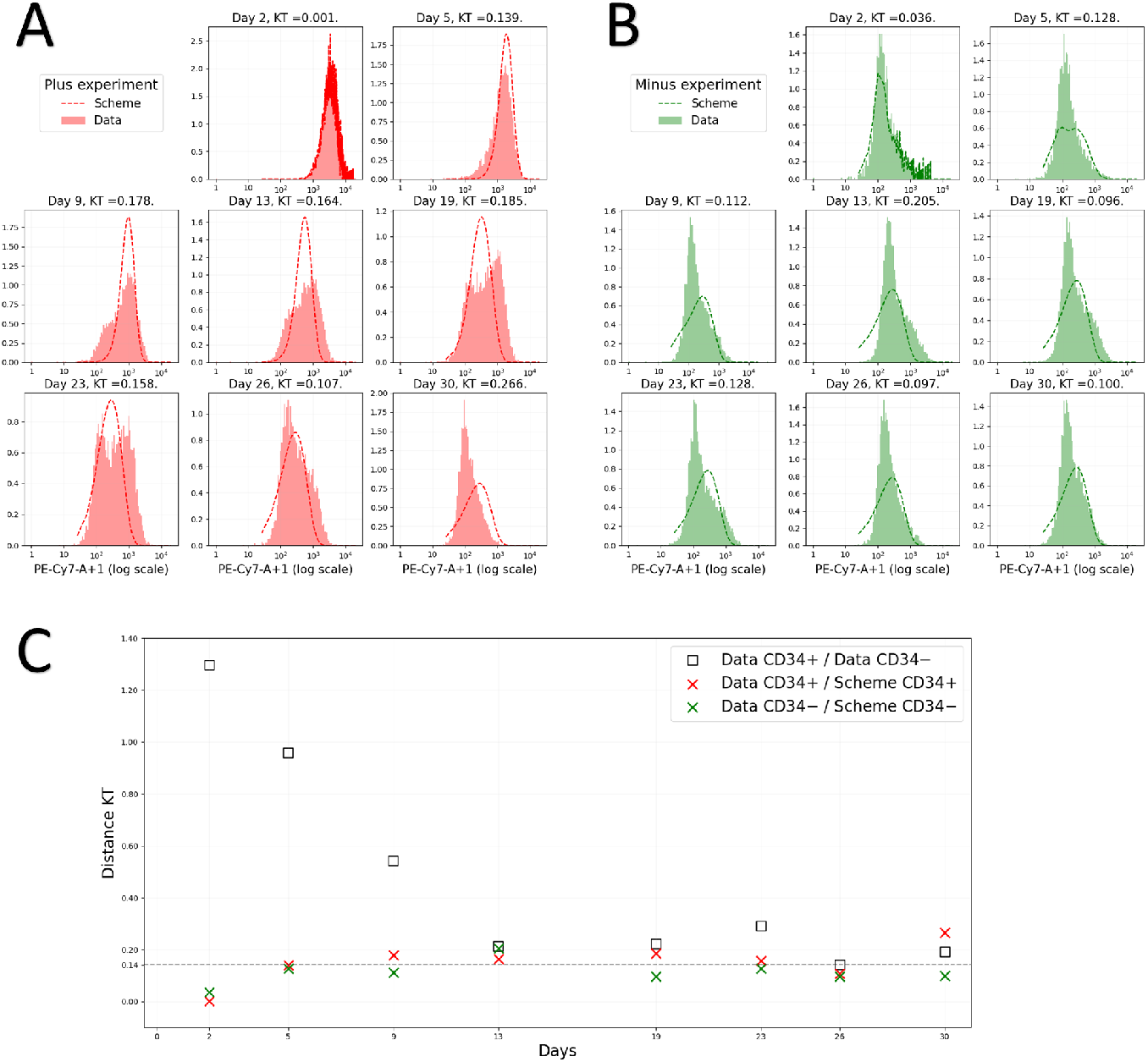
Comparison of model and data. On the left the fitting of the CD34^+^ relaxation experiment (in A) and on the right in green of the CD34^−^ (in B) experiments. Experimental data in logarithmic scale are represented by plain histograms and the numerical results of model (5) are represented by the dotted curves. We initialize the model on day 2, using the biological data. The initial condition is given by (24). Parameter values are given in table 1. KT: the modified Kantorovich-Rubinstein distance, defined by the equation (14). C. Time-dependent evolution of the Kantorovich-Rubinstein distance between model and experimental data. For different days of the experiment, the modified Kantorovich-Rubinstein distance between the two relaxation experiments is depicted using black squares. The minimum distance, reached on day 26, is illustrated by a horizontal dotted line. The red crosses correspond to the modified Kantorovich-Rubinstein distance between the model for the parameter values from Table 1, and the CD34^+^ cell relaxation experiment. Similarly, the green crosses represent the distance for the CD34^−^ cell relaxation experiment.

To initialize our model on day 2, we use the equation (8), it follows this following initial conditions

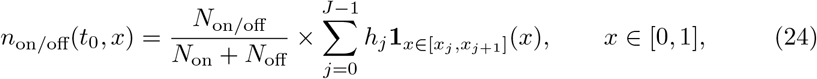

where *h*_*j*_ corresponds to the number of cells on day 2 with fluorescence between *X*_max_ × *x*_*j*_ and *X*_max_ × *x*_*j*+1_, where (*x*_*j*_)_*j*_ corresponds to the uniform discretization of space [0, 1]. That is, the (*h*_*j*_)_*j*_ correspond to the heights of the histogram of the data renormalized by the maximum *X*_max_.

Visually, Figure 6.A reveals a high degree of proximity between the experimental data and the mathematical model. To quantify this closeness, we once again employ the modified Kantorovich-Rubinstein distance. In figure 6.B, we represent, by black squares, the temporal evolution of distance between the experimental data of the two relaxation experiments. Due to the antinomic nature of the two experiments, distances are considerable in the early days of the experiment. It then gradually decreases as the two distributions converge towards the stationary distribution. From day 26, both profiles reached the stationary stage. At this point, in the absence of noise, these two distributions are expected to be similar. Therefore, the minimum distance, dist = 0.142 (illustrated by a dotted line), attained on day 26, corresponds to a reference distance to determine the proximity of two distributions.

In this figure, we also illustrate the distance between the model and the two relaxation experiments. The CD34^+^ cell relaxation experiment is represented by the red crosses, and the CD34^−^ cell relaxation experiment by the green crosses. To quantitatively assess the proximity of the model to experimental data, we employ the reference distance represented by the horizontal line. For the CD34^−^ cell relaxation experiment, we observe that the distances are always less than the reference distance, except on day 13 for which the distance is slightly greater. Concerning the CD34^+^ cell relaxation experiment, this time the distances are more regularly greater than the reference distance, but are still within an acceptable order of magnitude. These results show that our proposed model is very close to the experimental data.

## Discussion

In the present work, we proposed a revised two-state probabilistic model for gene expression which explicitly incorporates a proliferation term. This model was analysed and we obtained an analytical solution for our model’s steady state. The same model was then used for simulating the transient behavior of FACS-sorted cells leading to the progressive relaxation towards the steady state distribution. Altogether, our work shows that a two-state description for CD34 gene expression is well suited to explain the relaxation experiments.

Although we had to infer a number of model parameters which could not be deduced from the literature (like for example the half-life of the CD34 protein), the overall fitting ability of our model proved to be quite satisfactory. Using Kantorovich distances, we indeed observed that the model-to-experiment distance was within the range of the experiment-to-experiment distance, so in the range of experimental variability.

We assumed that the proliferation rate would depend upon the level of expression of the very gene that is being modelled. In our case, that proved to be useful since we wanted to fit relaxation data obtained from CD34 expression. CD34 is a known marker for stemness and we hypothesized that, in line with the existing litterature [22], CD34^+^ cells would proliferate less that CD34^−^ more mature cells. We should nevertheless stress that such a behaviour can be true for normal hemotopoietic stem cells, but can be questioned regarding cancer stem cells.

One of the difficulties we faced when comparing the model’s output with experimental data, lies in the need for common units By default, our model output is a value between 0 (no CD34 expressed) and 1 (maximum level of CD34 expression). FACS data are corrected fluorescent values, that can be negative in the raw acquisition dataset. We therefore processed the data with a gating phase, a shifting phase, and finally normalized them in order to obtain comparable values with the model.

It is crucial to emphasize that within a cell population displaying a stationary distribution of phenotypic states, no cell remains in a permanent state over time. Given a sufficiently long time, one can assume that all cells will have visited all possible states (i.e. all possible values for their surface CD34 expression). In other terms, in the state versus identity long standing debate [28], we clearly side with the view that stemness is an emerging dynamical property Several points shall be investigated further. The first point that can be enriched is the form of the division rate *r*(*x*) = *r*_0_ − *r*_1_*x*. The linear form ensures the explicit formulation of the stable distribution and facilitates the scaling by *X*_*max*_ but is not necessary for the existence of a profile. Moreover, we made strong assumptions here that the daughter cells have the same concentration of markers than the mother cell and that the division has no impact on the on*/*off status of the cell. This later is a reasonable assumption in the light of the existence of transcriptional memory [17, 18], but it might be gene-dependent.

One missing aspect of our model is the absence of any explicit death term. On the other hand, an expression independent death rate could immediately be considered by relaxation of the constraint of positive division rate (which would then correspond to a net growth rate). In terms of parameters, this would affect *r*_0_.

Another missing aspect of our model is the fact that the CD34 gene expression is modelled in isolation. It is quite obvious that in cells its expression level will be constrained by its positioning in a complex web of genes-to-genes interactions known as a Gene Regulatory Network (GRN). Inference of such GRNs is a notoriously difficult task (see e.g. [29]), and performing relaxation experiments from such complex objects is yet to be done.

One of the future goal of our work would be to assess its predictive ability. A promising lead would be to go further in the analysis in order to estimate the influence of parameters on the relaxation time. Mathematically, this could be analysed through the spectral gap which is beyond the scope of this work. It would be especially interesting to identify the effects of various parameters on it, in particular the parameter *d* which represents the degradation rate. Note that in this case, the distribution and the value of *λ* are expected to change. Interestingly, the model’s prediction in this case could be tested experimentally by modifying the endogenous CD34 protein stability.

## Supplementary Information

## Acknowledgments

We thank Ulysse Herbach for his help with inferring 2-states parameter values. We also thank the BioSyL Federation and the LabEx Ecofect (ANR-11-LABX-0048) of the University of Lyon for inspiring scientific events.

## Authors’ contributions

ME, OG, TL wrote the manuscript. ME, OG, TL and GR performed the mathematical derivation of the profile. MD and ME performed the numerical simulations analyzed the data. ME performed the parameter inference. ME, OG and TL interpreted the results. VMG, SL and TV produced the data. All authors but TV read and approved the final manuscript.

## Funding

Maxime Estavoyer is funded by ANR PLUME (ANR-21-CE13-0040). For this project, Marion Dufeu was funded by IXXI (http://www.ixxi.fr)

## Declarations

### Ethics approval and consent to participate

Not applicable

### Consent for publication

Not applicable

### Availability of data and materials

The datasets used and/or analysed during the current study are available from the corresponding author on reasonable request.

### Competing interests

## Appendix A Supplementary Figures

**Fig. A1.**
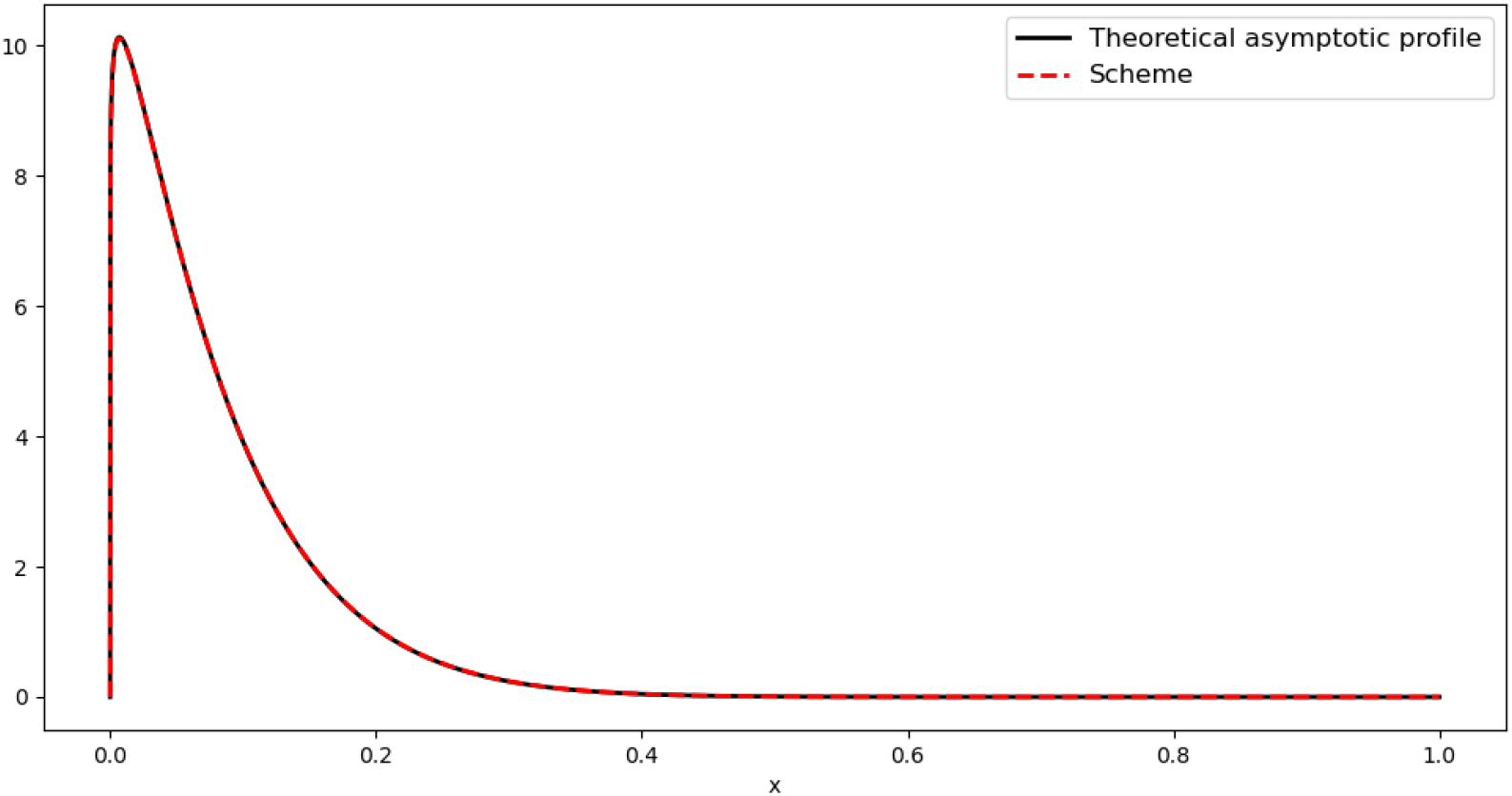
Comparison between the theoretical asymptotic profile *N* (black line) presented in Theorem 1 and the numerical solution obtained using the numerical scheme (red dotted line) defined by equations (9). Parameter values are given by *r*_0_ = 3.4, *r*_1_ = 3, *k*_on_ = 0.85, *k*_off_ = 12.71, *d* = 1, *X*_max_ = 2 *×* 10^4^.

## Notes

### Competing Interest Statement

The authors have declared no competing interest.

